# Caspase cleavage of APP contributes to amyloid beta-protein induced synaptic injury

**DOI:** 10.1101/2025.04.30.651606

**Authors:** Brea Midthune, Goonho Park, Sheue-Houy Tyan, Edward H. Koo

**Affiliations:** Department of Neurosciences, University of California, San Diego, La Jolla, CA; VA Palo Alto Healthcare System, Palo Alto, CA, USA; Departments of Medicine and Physiology, National University of Singapore Yong Loo Lin School of Medicine, Singapore

**Keywords:** Alzheimer’s disease, amyloid beta, amyloid-precursor protein, caspase, caspase-3, long-term potentiation, neurodegeneration, spine, synapse, synaptic depression

## Abstract

**BACKGROUND:** Increasing evidence suggests that amyloid beta (Aβ) lies at the center of Alzheimer’s Disease (AD) pathology and that synapses are the initial site of damage by Aβ. Recent studies have also indicated a role for caspases in AD-related synaptic dysfunction and memory loss, but the mechanism(s) through which the caspases act remains elusive. Previous studies in cell culture indicate that cleavage of a caspase site on the intracellular domain of the amyloid precursor protein (APP) protein contributes to Aβ−induced cell death. However, the role of this cleavage event in synaptic dysfunction has not been established.

**METHODS:** Through a combination of intracellular and extracellular electrophysiological methods and confocal microscopy of dendritic spines, we examined the involvement of caspase-3 and amyloid-precursor protein in Aβ-mediated synaptic dysfunction.

**RESULTS:** Here, we provide evidence that caspase activity at the intracellular domain of APP is required for acute Aβ-induced depression of glutamatergic synapses. We find that local elevation of Aβ levels through over-expression of the C-terminal fragment of APP (C99) failed to depress synapses if caspases were inhibited pharmacologically or in tissue lacking caspase-3. To demonstrate a link between these findings to APP, we found that Aβ failed to depress synaptic transmission or inhibit synaptic plasticity in neurons lacking APP. To specifically test the role of caspase cleavage of the intracellular domain of APP, we introduced a mutation that inhibits caspase cleavage at site 664 to the C99 construct; this construct produced Aβ but failed to elicit Aβ-induced synaptic depression or spine loss, and reduced caspase-3 activity.

**CONCLUSION:** Taken together, these results suggest an APP-dependent pathway in which caspases contribute to Aβ-induced synaptic depression and spine loss via cleavage of APP.

## Introduction

Alzheimer’s disease (AD) is a progressive neurodegenerative disorder that affects memory and cognition. Increasing evidence implicates amyloid beta (Aβ), a proteolytic product of amyloid precursor protein (APP), as the primary etiological factor and the synapse is thought to be the initial site of damage (Selkoe and Schenk 2003, Palop and Mucke 2010, Selkoe and Hardy 2016). APP is a type I transmembrane protein that is sequentially cleaved to generate multiple cleavage products, one of which is the toxic Aβ species. Although a number of cellular changes have been described secondary to Aβ-induced toxicity (Haass and Selkoe 2007, Benilova et al. 2012), the precise downstream mechanisms, especially in the *in vivo* setting, remain largely unresolved.

Consistent with the progressive neuron loss seen in late-stage AD, increased caspase activation is also detected in the brains of individuals with AD (Su et al. 1994, Yang et al. 1998, Gervais et al. 1999, Lu et al. 2000), with reports that the primary site of increased caspase activation is localized to synapses (Louneva et al. 2008). Caspase activation was reported to be upregulated early in AD progression (Gastard et al. 2003, Cribbs et al. 2004) and recently it was shown that increased caspase-3 activation is detectable in the absence of cell death in one APP transgenic mouse line (D’Amelio et al. 2011).

Interestingly, several studies have shown that caspase cleavage of APP contributes to Aβ-mediated cytotoxicity (Rohn et al. 2000, Bredesen et al. 2010). Caspase cleavage of APP at position 664 (using APP695 numbering) generates two major peptides: J-casp (from intramembrane γ-secretase cleavage site to APP664) and C31 (the C-terminal 31 amino acid peptide of APP from APP665), and, in cell culture, cytotoxicity has been attributed to both cleavage products (Lu et al. 2000, McPhie et al. 2001, Madeira et al. 2005, Shaked et al. 2006, Park et al. 2009). This view is supported by *in vivo* data that indicate increased caspase cleavage of APP in brains of individuals with AD correlates with disease severity (Gervais et al. 1999, Lu et al. 2000, Banwait et al. 2008). Additionally, preventing caspase cleavage of APP by mutating the critical aspartate residue at APP position 664 has been associated with a reduction in amyloid-associated pathology in transgenic mice, including reduced synapse loss or impairment in synaptic plasticity but without any reduction in Aβ levels (Galvan et al. 2006, Saganich et al. 2006), although the results were debated in a later study (Harris et al. 2010). Thus, the precise pathological contribution of caspase cleavage of APP in Aβ-induced synaptic dysfunction in the absence of cell death remains unresolved (Bredesen et al. 2010).

Here, we demonstrate that Aβ synaptic toxicity can, in part, be attributed to caspase cleavage of APP in an organotypic slice culture (OTSC) system. Caspase activation was associated with localized synaptic injury as assessed by Aβ-mediated depression of neurotransmission and by reduction of dendritic spines. Further, the presence of both APP and an intact caspase cleavage site in APP are necessary for Aβ-mediated synaptic depression and LTP impairment. These changes correlated with the presence of increased caspase-3 activity in dendrites and spines.

## Experimental methods

### Transgenic Mice

Caspase-3 KO animals (line B6.129S1-*Casp3tm1Flv*/J) were obtained from Jackson Laboratories. APP KO and APLP2 KO mice were kindly provided by Dr. Hui Zheng at Baylor College of Medicine (Zheng et al. 1995, von Koch et al. 1997). All transgenic mouse lines were maintained in a C57BL/6 inbred background.

### Organotypic Slice Culture

Hippocampal slice cultures were prepared from 5- to 8-day-old Sprague Dawley rats or transgenic mice, essentially as described previously (Stoppini et al. 1991). Briefly, hippocampal slices were cut in ice cold low sodium cutting buffer at a thickness of 400 μM using a McIIwan Chopper and maintained on semipermable membrane inserts and maintained at 35° C. Culture medium was changed every 2-3 days.

### Constructs and Virus Preparation

Sindbis virus containing APP β-CTF (C99) IRES eGFP, C99 IRES tomato or tomato only were used as described (Kamenetz et al. 2003). An aspartate to alanine substitution was introduced at APP site 664 (D664A) of the C99 IRES eGFP or tomato construct using site-directed mutagenesis (Quikchange, Stratagene). Sindbis virus was prepared using an expression system (Invitrogen) and in accordance with manufacturer’s directions. Virus was injected into the CA1 region of hippocampal slices to test for infectivity as demonstrated by fluorescent protein expression and diluted to the desired concentration.

### Intracellular Electrophysiology of Organotypic Slice Culture

Organotypic hippocampal slices were locally injected using a glass pipette and picospritzer to sparsely infect CA1 neurons with Sindbis virus expressing the desired protein and GFP (or GFP alone, as control) at day 6-8 DIV and whole cell patch recordings of CA1 neurons were performed at least 14 hours after infection (Kamenetz et al. 2003). Whole cell recordings were done in pairs as previously described (Kamenetz et al. 2003). Results from two pathways recorded at the same time were averaged and counted as *n* = 1. Pairs were counted if the input resistance was at least 10 fold higher than the series resistance. For caspase inhibitor experiments, slices were pre-incubated in medium containing either 100 μM of z-VAD-fmk, 100 μM z-FA-fmk, or 10 μM z-DEVD-fmk (R&D Systems) overnight. Equal parts DMSO were used as negative control and experiments were done blind to the experimenter.

### Immunoassays

Antibodies, including procaspase 3, cleaved caspase 3, and β-actin, were purchased from Cell Signaling and Sigma. As briefly explained, protein samples from OTSC were extracted by 1X RIPA buffer containing 1X protein inhibitor (Sigma-Aldrich) and the protein concentration was measured by Biorad protein assay (DC^TM^ protein assay, #5000111). 20ug protein was run on 10% Bis/Tris gel and transferred nitrocellulose membrane. Antibodies used: CT-15 (1:10,000) for full length of APP; 6E10 (1:1000) for C99; procaspase 3 (1:1000) and cleaved caspase 3 (1:1000) for caspase 3; β-actin (1:1000) for loading control.

### Analysis of spine loss after C99 and C99_D664A expression

The Sindbis viral vectors expressing Tomato (control), C99_wild type (WT) or C99_D664A mutant co-expressing fluorescent protein were injected into 5-day-old OTSCs from wild-type mice. After infection, the slices were incubated for 1 day without changing media. The OTSC tissues were fixed with fixing solution (4% PFA/ 3% sucrose) for 30 minutes and washed with 0.1% PBS-T solution and mounted with mounting solution (Vector Shield, #H1200). To analyze synaptic density and caspase activation, we used 0.5μM ultra bright green dye-DEVD-fmk, a fluorochrome inhibitor of caspases (FLICA) (Vergent Bioscience, # 13101) to detect and image caspase activation in live dendritic spines using the manufacturer’s protocol. Confocal microscopy (Olympus FV1000 Confocal Microscopy, 100X oil objective) was used to analyze alteration of synaptic density induced by sindbis viral infection.

### Preparation for 7PA2-conditioned medium

Conditioned medium was collected from Chinese hamster ovary (CHO) cells stably expressing human APP751 with the Val717Phe mutation (7PA2 cells) (Podlisny et al. 1995) that were cultured in DMEM with10% fetal bovine serum as described previously (Walsh et al. 2002, Calabrese et al. 2007). Cells were grown to near confluence and washed with PBS, and then maintained in plain DMEM for ∼16 h. The 7PA2 conditioned medium (7PA2-CM) was concentrated ∼10-fold using YM-3 Centriprep filters (Amicon, Beverly). Concentration of Aβ42 was determined by ELISA assay and stored at -80°C. The ELISA utilized mAb Aβ 42.2 (Levites et al. 2006) (kindly provided by Todd Golde, UFL) as capture and HRP-conjugated 6E10 (Covance,) as reporter.

### Acute hippocampal slice electrophysiology

Acute coronal hippocampal slices and LTP were performed as described (Midthune et al. 2012) from 2-4 month old APP or APLP2 KO mice and their control littermates. Amplitudes were measured and compared at 50-60 minutes post-tetanus.

### Statistical analysis

Immunoblotting results and images of caspase activation were quantified using ImageJ. Data of immunoblotting, caspase assay, and Aβ ELISA experiments were analyzed with Student’s t-test and one-way ANOVA using GraphPad. LTP data were analyzed using the unpaired Student’s t-test, while paired intracellular recordings of were analyzed using the paired Student’s t-test. The AMPA/NMDA ratios were analyzed using the unpaired Student’s t-test or one-way ANOVA. All error bars indicate standard error of mean (SEM). A p-value less than 0.05 was considered statistically significant.

## Results

### Overexpression of C99 induces Aβ-mediated synaptic depression

First, we tested the functionality of our system for measuring synaptic depression in neurons overexpressing Aβ. This system allows for the direct comparison of post-synaptic transmission in adjacent neurons (Kamenetz et al. 2003). Here, we sparsely infected neurons in the CA1 hippocampal region with Sindbis virus expressing APP C-terminal fragment, C99, which was constructed with an IRES eGFP sequence to identify infected neurons. Evoked synaptic currents of cells overexpressing C99 were compared to adjacent uninfected neurons using dual whole-cell path clamp recordings as previously described (Kamenetz et al. 2003) (Fig. 1A & B). Consistent with previous reports, overexpression of C99 led to robust depression of α-amino-3-hydroxy-5-methyl-4-isoxazolepropionic acid receptor-(AMPAR) and N-methyl-D-aspartate receptor- (NMDAR) (but not gamma-aminobutyric acid receptor- [GABAR]) mediated excitatory post-synaptic currents (EPSCs) (Shi et al. 2001, Kamenetz et al. 2003, Hsieh et al. 2006) (Fig. 1C). To test for changes in basal synaptic strength through alterations in relative numbers of AMPA and NMDA receptors, the AMPA/NMDA ratio was calculated and it remained unchanged in neurons expressing C99 (Figure 1D). Importantly, we know the resulting synaptic depression is specific to the overexpression of C99 and is Aβ dependent, as this has been demonstrated previously by Kamenetz et al. (2003) and others (Hsieh et al. 2006, Kessels et al. 2013).

**Figure 1:**
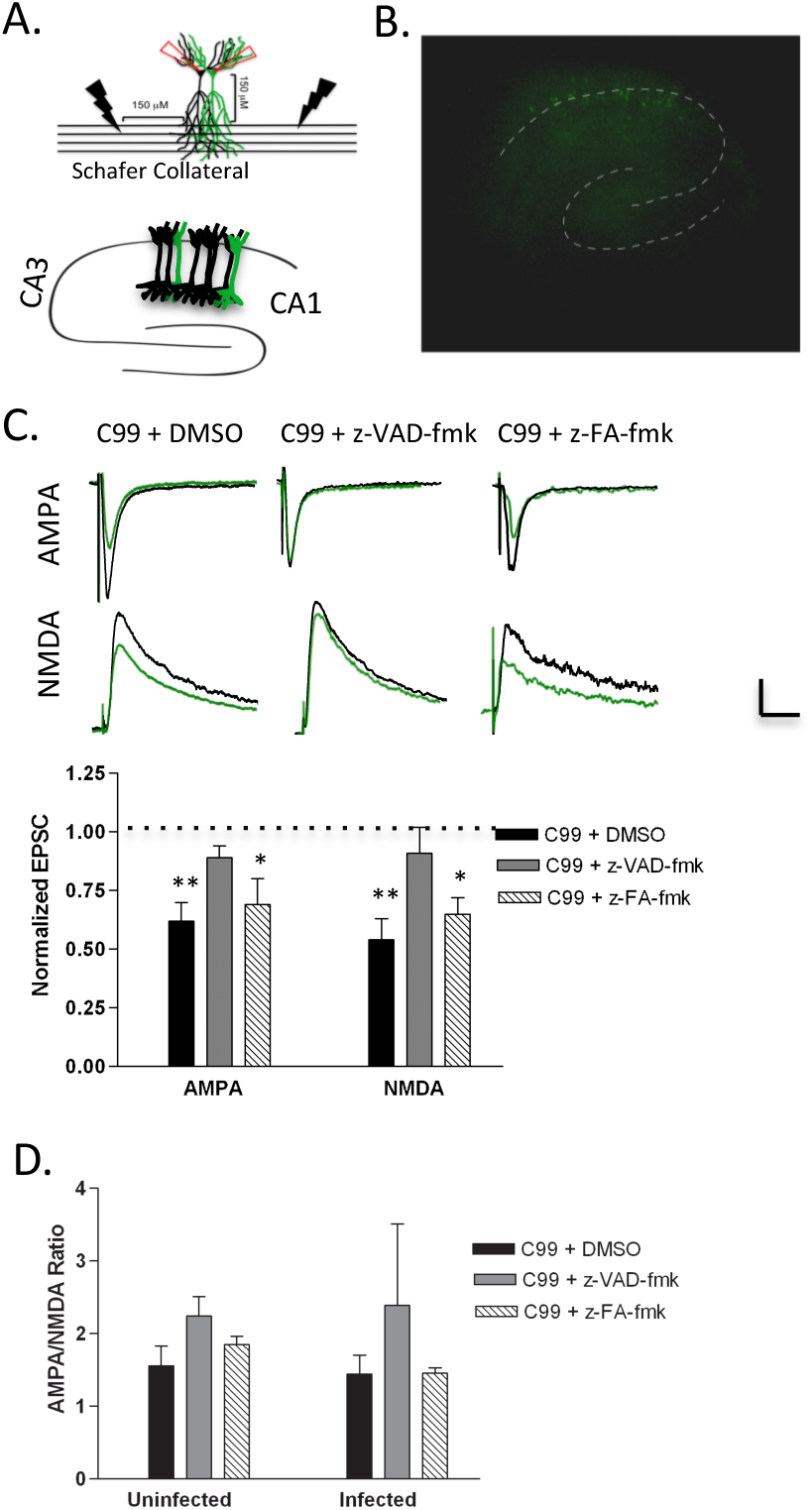
A) Top, Schematic of paired, whole cell recording method. EPSCs are recorded simultaneously from neuronal pairs and evoked with bipolar simulating electrodes, allowing for direct comparison of synaptic currents. Bottom, CA1 pyramidal neurons were sparsely infected with sindbis virus expressing APP β-CTF (C99) IRES eGFP. B) A representative image of a hippocampal slice sparsely infected with virus producing C99 IRES eGFP. C) Top, Average of 30-60 sweeps recorded simultaneously from infected (expressing C99, green) and non-infected (black) neurons from slices treated as indicated. Bottom, Graph of the average of synaptic responses in infected neurons normalized to the average of response in non-infected neurons for sparsely infected slices exposed to indicated drugs (C99+: DMSO, AMPA: p=0.002, n=11 pairs; NMDA: p=0.042, n=11; caspase inhibitor z-VAD-fmk, AMPA: p=0.19, n=13; NMDA: p=0.5, n=10; inhibitor control z-FA-fmk, AMPA: p=0.013, n=13; NMDA: p=0.025, n=9). D) The AMPA to NMDA ratios were compared across conditions using single factor ANOVA and showed no significant difference between conditions for uninfected or infected neurons. Uninfected: F(2,32)=0.483, p=0.621, Infected: F(2,31)=0.844, p=0.440.

### Inhibition of caspases blocks Aβ-mediated synaptic depression

Many studies have suggested that caspases play a role in Aβ-mediated neurodegeneration (reviewed in Rohn and Head 2009). Additionally, recent evidence indicates that caspases are localized in the synapse (under both physiological and pathological conditions) and play an important role in synaptic function (Li et al. 2010, Erturk et al. 2014, Shen et al. 2014). However, whether caspases contribute specifically to Aβ-mediated synaptic dysfunction, especially in acute toxicity, is less defined. To test the latter hypothesis, OTSCs sparsely infected with Sindbis virus overexpressing C99, were incubated overnight with z-VAD-fmk, a cell-permeant pan-caspase inhibitor, or z-FA-fmk, a well-characterized control for fmk conjugated caspase inhibitors. We found that addition of z-VAD-fmk blocked synaptic depression in neurons overexpressing C99 (Fig. 1C), while z-FA-fmk had no effect (Fig. 1C), supporting the concept that caspases contribute to Aβ-mediated synaptic depression. To determine whether z-VAD-fmk or z-FA-fmk affects basal synaptic strength, we analyzed AMPA/NMDA ratio and found no significant difference between conditions (Fig. 1D).

Because prior studies have implicated caspase-3 as an important effector caspase in Aβ toxicity and synaptic plasticity, the OTSCs were incubated with the caspase-3 inhibitor, z-DEVD-fmk (Li et al. 2010, D’Amelio et al. 2011, Jo et al. 2011). Treatment with z-DEVD-fmk restored AMPA and NMDA transmission to normal levels in neurons overexpressing C99 (Fig. 2A). To further support this result, we then confirmed the role of caspase-3 in Aβ-mediated synaptic depression by testing the effect of C99 overexpression in caspase-3 knockout mice. Consistent with our previous results, we found a similar attenuation of Aβ-mediated synaptic depression (Fig. 1D). Again, to account for the possibility that basal synaptic strength may be altered by caspase-3 inhibition or deletion, we analyzed AMPA/NMDA ratio and found no significant changes (Fig. 2B). Together, both pharmacologic and genetic targeting support the view that caspases, in particular caspase-3, contribute to Aβ-mediated impairment of glutamatergic neurotransmission.

**Figure 2:**
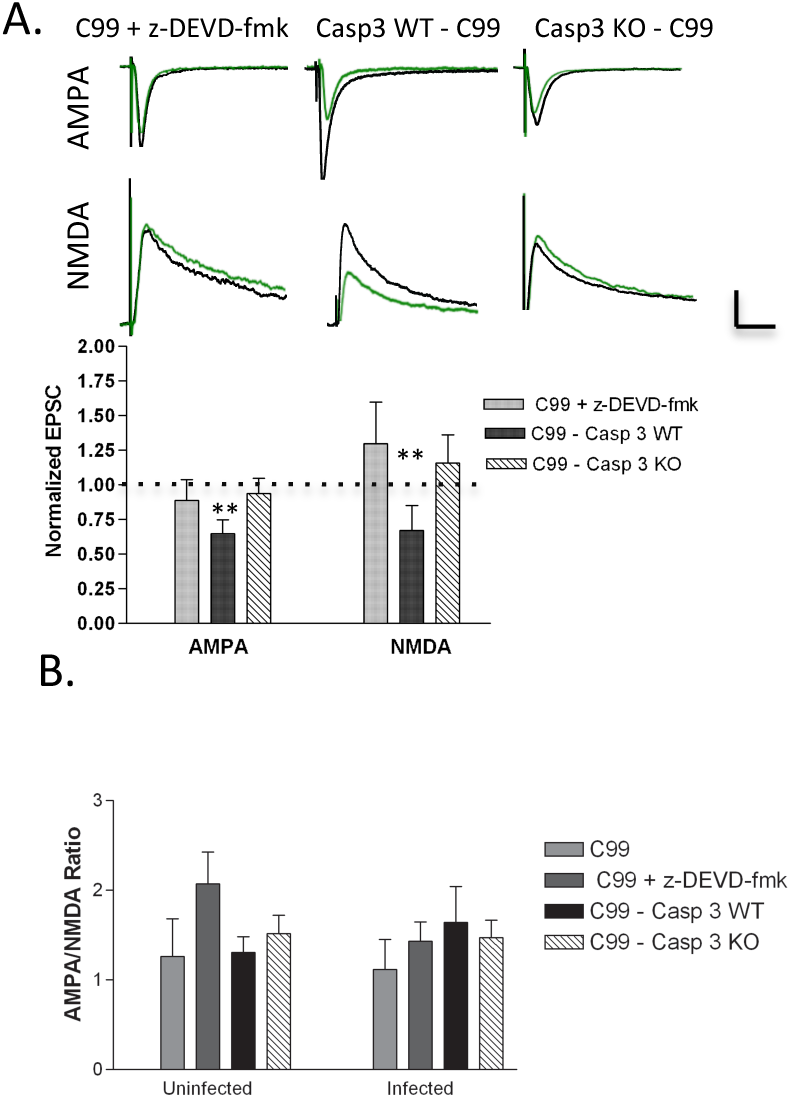
Treatment with caspase inhibitors prevented Aβ-induced depression of AMPAR-mediated synaptic currents. A) C99 expressed with caspase-3 inhibitor z-DEVD-fmk, (AMPA: p=0.57, n=10; NMDA: p=0.398, n=9); C99 expressed in a caspase-3 WT background (AMPA: p=0.001, n=12; NMDA: p=0.008, n=11); C99 expressed in a caspase 3 KO background (AMPA: p=0.59, n=14; NMDA: p=0.323, n=13). Dotted line represents the normalized mean amplitude of the uninfected control cell. Error bars indicate mean ± SEM. *p <0.05, **p <0.01. Scale bars, 60 pA, 40 ms. B) The AMPA to NMDA ratios showed no significant difference between conditions. AMPA/NMDA ratio (uninfected): F (3,37) = 1.290, p=0.292; AMPA/NMDA ratio (infected): F (3,37) = 0.099, (p=0.960)

### Deletion of APP attenuates Aβ-mediated synaptic depression

As documented here and by others, the presence of caspase-3 is necessary for Aβ-mediated synaptic depression (D’Amelio et al. 2011, Jo et al. 2011). However, it is unclear which of the many caspase substrates contribute to this pathway of Aβ toxicity upon caspase activation. Because our previous studies showed Aβ-mediated cell death exhibited an APP-dependent mechanism in cultured cells (Lu et al. 2003, Shaked et al. 2006, Park et al. 2009), we questioned whether APP contributes to this experimental paradigm of Aβ-mediated synaptic depression. We tested this hypothesis by comparing the effects of C99 overexpression in wild type and APP deficient animals. In these experiments we used a dense, rather than sparse, infection protocol. This dense infection produces mass amounts of extracellular Aβ, resulting in synaptic depression in both infected and nearby non-infected neurons by diffusion of secreted Aβ (Kamenetz et al. 2003) (Fig. 3A). Therefore, if APP is required for Aβ-mediated synaptic depression, absence of APP should block Aβ-mediated synaptic depression, and the amplitude of AMPAR- and NMDAR-mediated EPSCs of the non-infected neurons should be increased relative to the C99 expressing neurons. As shown previously (Kamenetz et al. 2003), OTSCs from wild-type animals that were densely infected with C99-expressing Sindbus virus show similar NMDAR- and AMPAR-mediated responses between uninfected and infected neurons (Fig. 3B), indicating that both neurons undergo similar Aβ-mediated synaptic depression. In contrast, in OTSCs from APP KO animals, a dense infection led to depressed AMPAR-mediated EPSCs, but only in neurons infected with C99 (Fig. 3C), indicating that the C99 fragment of APP contributes to Aβ-mediated depression of AMPAR-mediated currents. Notably, we did not see a similar result for NMDAR-mediated EPSCs. Finally, to account for the possibility that basal synaptic strength may be altered by the absence of APP, we analyzed AMPA/NMDA ratio and found no significant change (Figure 3D). These results suggest that APP, and more specifically the C99 fragment of APP, contributes to Aβ-mediated synaptic depression.

**Figure 3:**
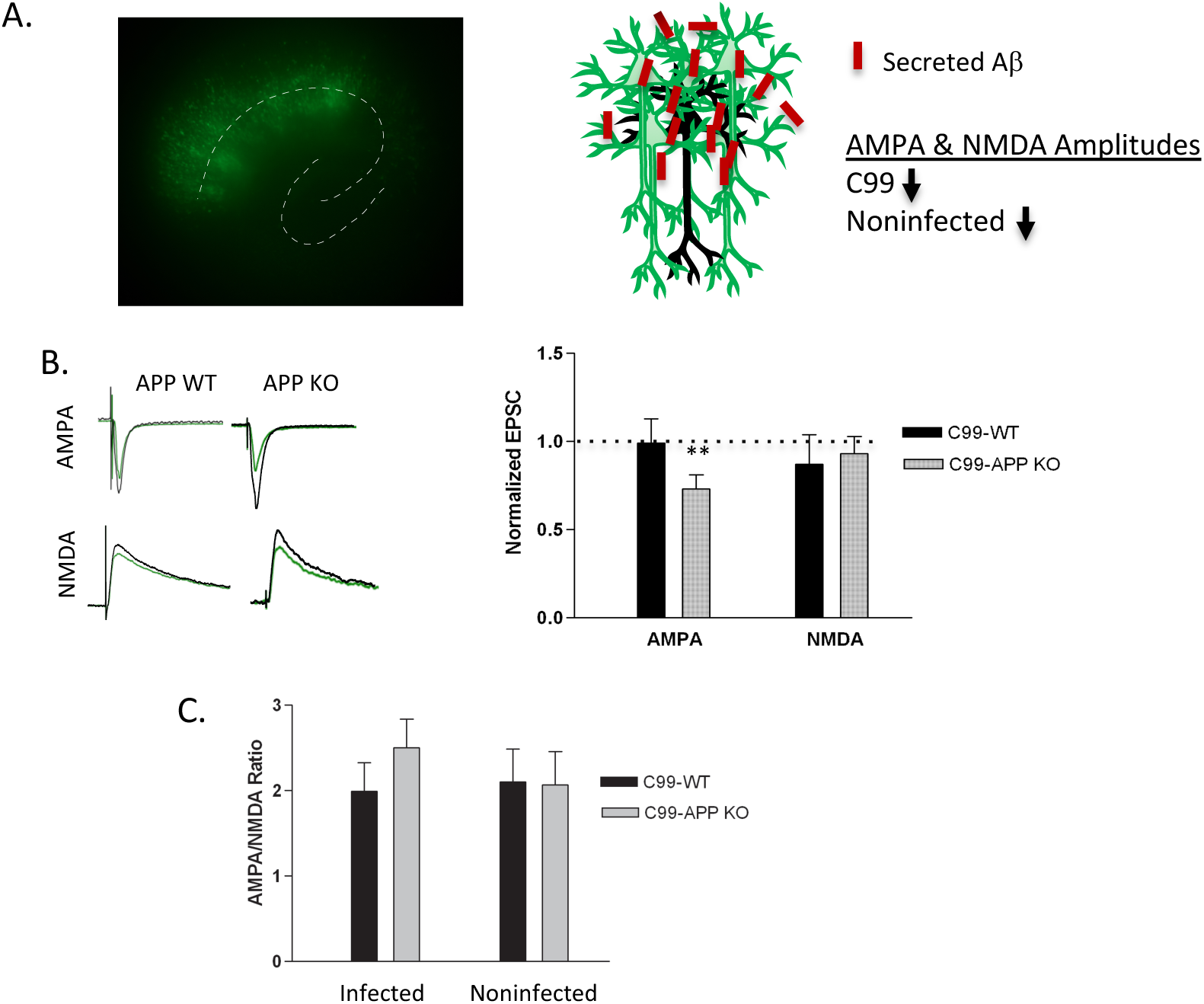
APP deficient neurons are protected from Aβ−induced reduction in AMPA R-mediated transmission. A) Left: A representative image of a hippocampal slice densely infected with virus producing C99 IRES eGFP. Right: Schema of experimental design. Because the dense infection produces mass quantities of secreted Aβ, both neurons overexpressing C99 and noninfected neurons experience synaptic depression. Averages of 30-60 sweeps recorded simultaneously from infected (expressing C99, green) and non-infected (black) neurons from slices obtained from indicated genotype. Graph of the average of synaptic responses in infected neurons normalized to the average responses in non-infected neurons for densely infected slices made from animals of indicated genotype (C99 expressed in APP WT, AMPA: p=0.91, n=12; NMDA: p=0.302, n=12; C99 expressed in APP KO, AMPA: p=0.006, n=14; NMDA: p=0.482, n=11). D) The AMPA to NMDA ratios showed no significant difference between conditions. AMPA/NMDA ratio (uninfected): p=0.24; AMPA/NMDA ratio (infected): p=0.95)

### Deletion of APP reduces Aβ-induced LTP impairment

Aβ has been shown to impair LTP (Walsh et al. 2002, Wang et al. 2004). To determine if APP is required for Aβ-induced impairment of LTP, we incubated acute brain slices from 3-4 month old APP KO or wild type mice in Aβ-conditioned 7PA2 medium (as source of Aβ oligomers) or control CHO medium and induced LTP with high frequency stimulation. While synaptic plasticity was reduced in slices from wild-type mice after Aβ incubation as expected (Fig. 4A), LTP was unimpaired in slices obtained from APP KO animals after Aβ incubation (Fig.4B), further supporting the view that APP is required for Aβ-induced synaptic dysfunction (Walsh et al. 2002, Wang et al. 2004). To test the specificity of APP in this pathway, we also examined slices from APLP2 KO animals. APLP2 is a close homologue of APP with high sequence identity. It shares similar processing and functional domains, but importantly, APLP2 does not contain the Aβ region, and there are differences in the intracellular terminal C31 sequence, which is one of two putatively cytotoxic peptides in APP that is released through caspase cleavage at the D664 site (Galvan et al. 2002, Eggert et al. 2004, Jacobsen and Iverfeldt 2009). Indeed, deletion of APLP2 did not restore LTP in the presence of Aβ (Fig. 4C), thus supporting the idea that APP contributes to Aβ-mediated depression of LTP.

**Figure 4:**
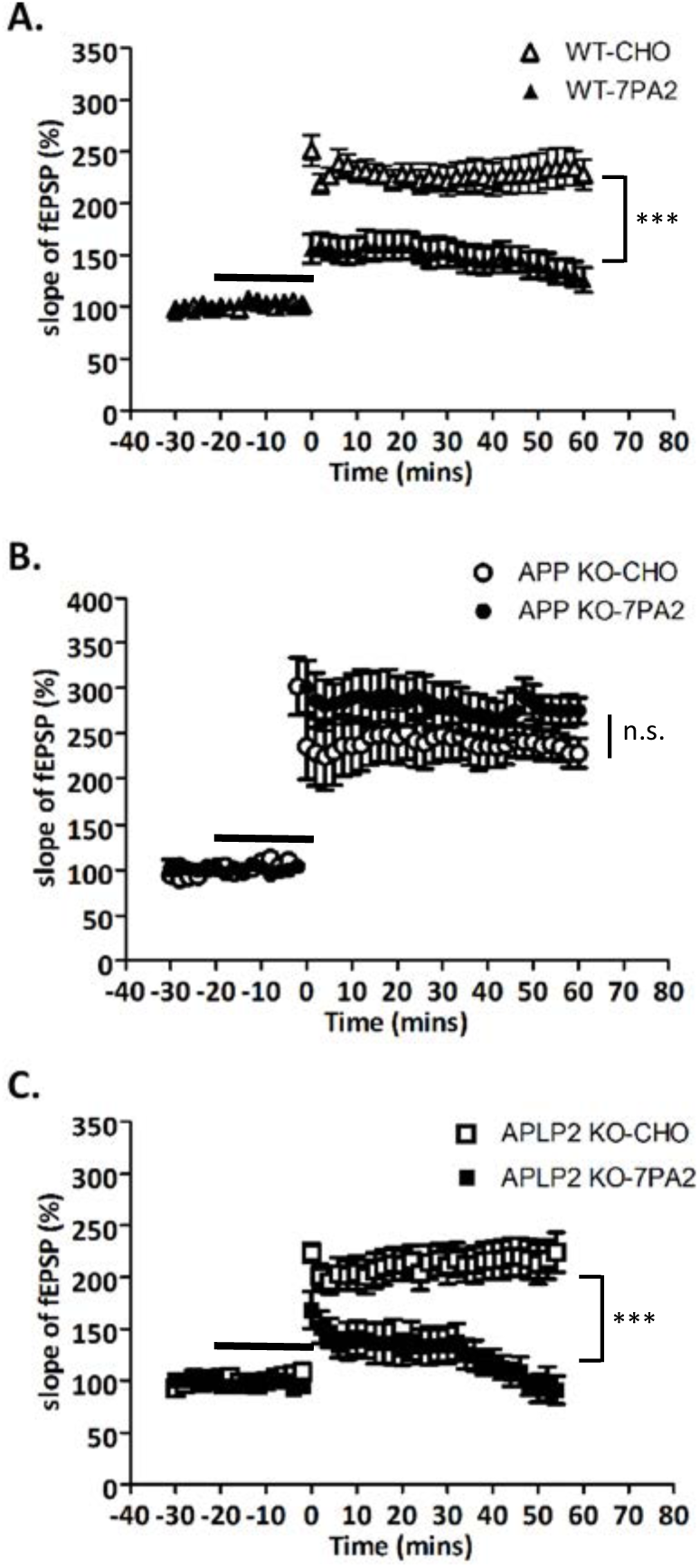
APP but not APLP2 deficiency attenuated Aβ-mediated LTP impairment. A) LTP in CA1 hippocampus from wild type (WT) mice after a 20-minute incubation in Aβ conditioned media (7PA2). Conditioned media from untransfected cells (CHO) were used as control (p=0.0001, CHO: n=7, 7PA2: n=8). B) LTP in APP deficient (APP-KO) mice after a 20 minute incubation in 7PA2 media or control CHO media (p=0.12, CHO: n=7, 7PA2: n=8). C) LTP in APLP2 deficient (APLP2-KO) mice after a 20 minute incubation in 7PA2 media or control CHO media (p=0.0004, CHO: n=5, 7PA2: n=6), ***p<0.001.

### Mutation of APP caspase cleavage site D664 prevents synaptic depression induced by Aβ

The data presented above indicate that the C99 fragment of APP, as well as caspase activity, are required for Aβ-mediated depression of AMPAR-mediated EPSCs and that APP is required for Aβ-mediated impairment of LTP. One mechanism potentially underlying these APP-dependent effects, thus linking the two mechanisms together, is the caspase-mediated cleavage of APP at position 664. To test if caspase cleavage of APP is indeed required for Aβ-induced synaptic depression, we overexpressed C99 with an aspartate to alanine mutation at site 664 (D664A), which prevents cleavage by caspases (Lu et al. 2000, Galvan et al. 2002). First, we confirmed that mutation at this position did not alter Aβ production, a finding consistent with previous reports (Fig. 5A, B, C) (Soriano et al. 2001, Tesco et al. 2003). Next, we found that the D664A mutation was protective against Aβ-induced synaptic depression (Fig. 5D), suggesting that the cleavage of APP by caspases contributes to Aβ-mediated synaptic toxicity.

**Figure 5:**
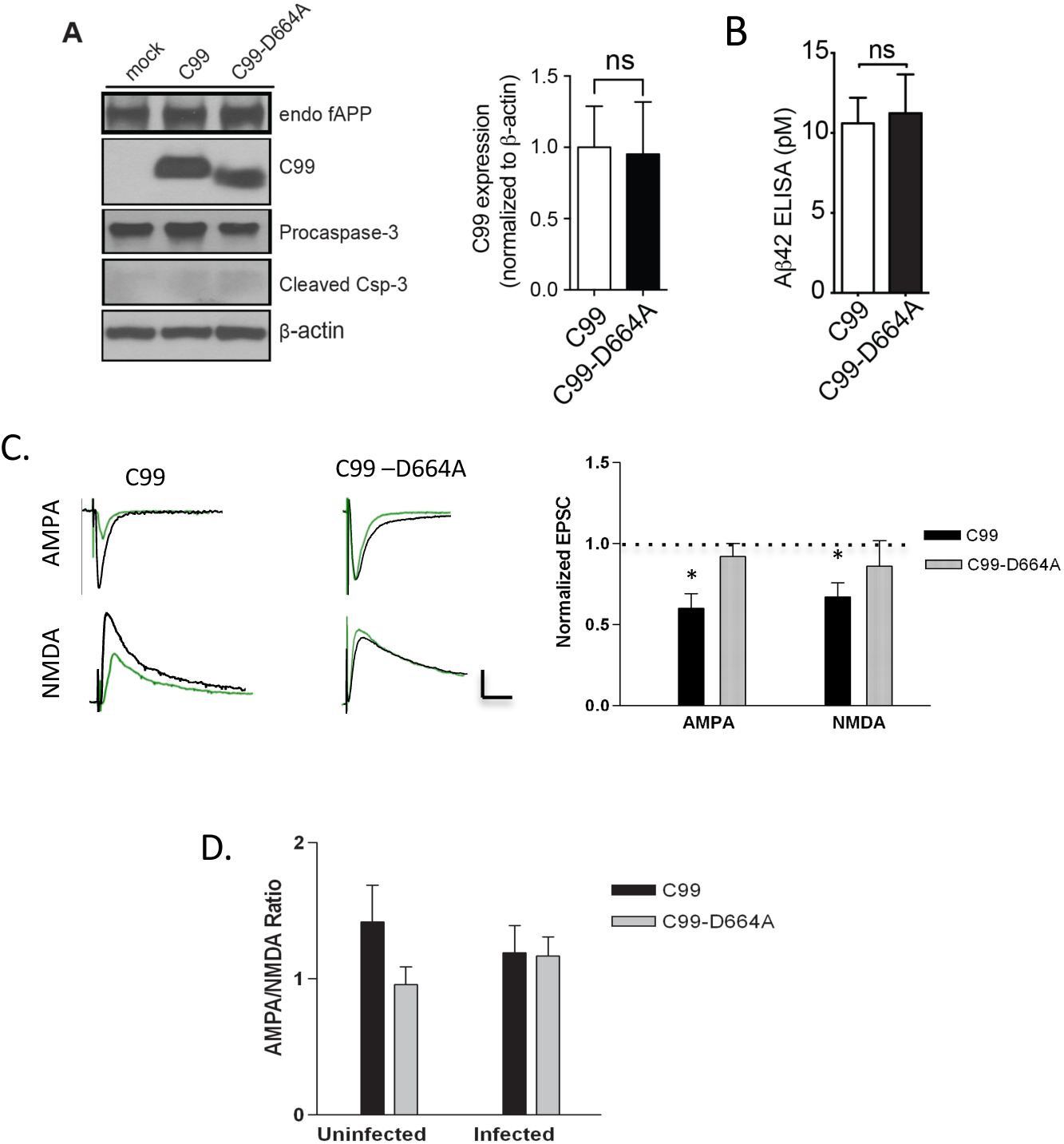
Elimination of internal APP caspase cleavage site prevents Aβ-mediated synaptic depression. A) C99_D664A produces similar levels of C99 compared to C99 WT, p=0.8225. B) C99_D664A produces similar levels of Aβ compared to C99 WT, p=0.965. C) Top, Averages of 30-60 sweeps were recorded simultaneously from infected (green) and non-infected (black) neurons expressing indicated constructs. Bottom, Graph of the average of synaptic responses in infected neurons normalized to the average of response in non-infected neuron for sparsely infected slices expressing indicated construct (C99, p=0.015, n=12; NMDA: 0.020, n=12; C99-D664A, AMPA: p=0.39, n=16; NMDA: p=0.219, n=15). D) The AMPA to NMDA ratios showed no significant difference between conditions: AMPA/NMDA ratio (uninfected), p=0.24; AMPA/NMDA ratio (infected), p=0.95.

### Mutation of APP caspase cleavage site reduces caspase-3 activity in neurons and attenuates Aβ−mediated synapse loss

To assess whether the Aβ and caspase-dependent reduction of NMDAR- and AMPAR-mediated EPSCs are accompanied by structural changes, OTSCs were sparsely infected with Sindbis virus expressing Tomato (control), C99 WT, or C99-D664A were assessed for synaptic changes and caspase activity by confocal microscopy. Indeed, in OTSCs infected overnight with Sindbis virus, there was ∼45% reduction in dendritic spine density in C99 WT but not C99 D664A infected neurons, which is consistent with our electrophysiological data. Furthermore, using a fluorogenic caspase reporter, we found that the D664A mutation significantly reduced caspase-3 activity in both dendrites and spines (Figure 6A). On the other hand, treatment of z-VAD-fmk could prevent Aβ-induced spine loss (Figure 6B). These results suggest that blocking APP C-terminus cleavage protects caspase-mediated spine loss.

**Figure 6:**
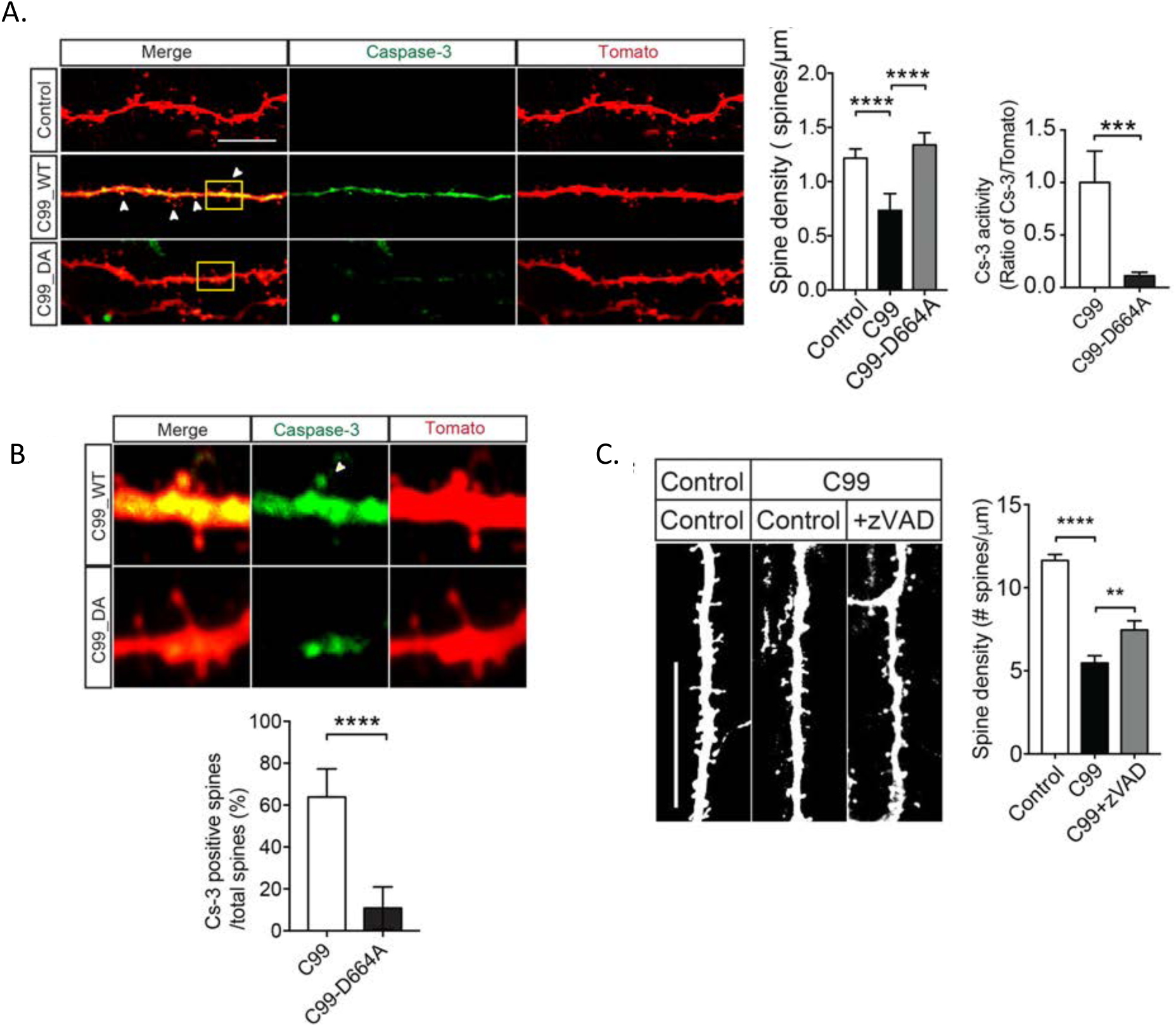
Elimination of internal APP caspase cleavage site and prevents spine loss and reduces caspase activity. A) Upper: Representative image of neurons infected with control-tomato, C99-tomato or C99-D664A-tomato and stained by DEVD fluorogenic reporter for caspase-3 activation. Scale bar 10 μm. Lower: Spine density (left) at 24hours after infection. Neurons infected with C99 exhibited a reduced number of spines compared to controls (p<0.0001) as well as increased caspase 3 staining (right). Neurons infected with C99-D664A recovered dramatically the spine loss compared with C99 WT (p<0.0001) and also showed decreased caspase activity compared to neurons infected with C99 (p=0.0009). B) Statistical analysis of dendritic spine density, which was infected with control and C99 WT and treated with 100μM z-VAD. Inhibition of caspase activation by pan-caspase inhibitor rescued C99-induced spine loss (C99 to control: ns, not significant, p=0.2403).

## Discussion

While caspases are essential in many cell death pathways, their role in neurodegenerative diseases, especially AD, is uncertain. Recent reports indicate that caspases may be important for neuronal processes extending beyond apoptosis, such as synaptic structure and function in both physiological and pathological contexts (Li et al. 2010, D’Amelio et al. 2011, Jo et al. 2011). In this study, we provide evidence that caspase activity is required for Aβ-induced synaptic dysfunction that acts, in part, through caspase cleavage of APP.

Caspase activity has been linked to Aβ-induced impairment of LTP (Jo et al. 2011), Aβ-mediated spine loss (Pozueta J 2013), and Aβ-mediated depression of basal glutamatergic synaptic transmission in an AD mouse model (D’Amelio et al. 2011) in the absence of overt cell death. We provide evidence that caspase activity is required in synaptic depression produced by acutely and locally delivered Aβ. We also used both pharmacological inhibition and genetic manipulation, as caspase inhibitors are well known to act off target and inhibit other caspases (Berger et al. 2006). Our results support these aforementioned reports by showing that caspase-3 activity contributes to depression of glutamatergic transmission mediated by Aβ.

While APP is a substrate to number of caspases *in vitro* (Galvan et al. 2002) it is unclear which caspases cleave APP *in vivo*. Though the reports by D’Amelio et al., (2011) and Jo et al., (2011) led us to focus on caspase-3, we cannot rule out the importance of other caspases. Notably, caspase-3 activity is dependent on upstream caspase activity, suggesting that other caspases must be involved (Salvesen and Riedl 2008).

Though initiating events of this upstream caspase activity are not addressed here, a series of reports have directly linked caspase activation to Aβ and APP oligomerization. Specifically, these studies showed that caspase activation is initiated through the binding of Aβ (Shaked et al. 2006, Lefort et al. 2012) to the cognate extracellular domain of APP (or C99) and causing APP oligomerization. The oligomerization has been shown to recruit caspases and induce caspase activation (Lu et al. 2003, Shaked et al. 2006) and toxicity (Rohn et al. 2000, Sudo et al. 2000) suggesting an APP-dependent component in Aβ-mediated cell death *in vitro*. Notably, a recent report showed human brain-derived Aβ oligomers bind to synapses and disrupt synaptic activity, including LTP and EPSCs, in a APP-dependent manner, though the mechanism of disruption was not tested (Wang et al. 2017).

Here, we found that reintroducing C99 into neurons lacking APP permits Aβ-mediated synaptic depression, consistent with the view that APP (and in our system, C99) may act as a receptor to Aβ, leading to synaptic depression of AMPAR-mediated EPSCs. Notably, reintroduction of C99 into neurons lacking APP does not permit depression of NMDAR-mediated EPSCs; this suggests that the Aβ-induced synaptic depression of NMDAR-mediated EPSCs are controlled by signaling mechanisms that differ from AMPAR-mediated EPSCs.

Interestingly, another group recently showed that the overexpression the APP intracellular domain affects synaptic function, including LTP and LTD, independent of Aβ toxicity. Their studies also found that caspase cleavage at the D664 site is responsible for this effect, in a model similar to ours (Trillaud-Doppia et al. 2016, Trillaud-Doppia and Boehm 2018), suggesting that caspase cleavage of APP may be important for synaptic function under physiological conditions. Here, we report that APP is required for Aβ-induced LTP deficits, specifically through caspase cleavage of APP at site D664. Furthermore, Aβ-dependent spine loss and caspase activation can be prevented by inhibition of APP C-terminus cleavage.

In summary, our data indicate that caspase activation, and in particular, cleavage of APP, contributes to acute Aβ-induced synaptic depression of AMPAR-mediated postsynaptic transmission, long-term potentiation, and spine loss. The mechanisms of toxicity through this caspase cleavage event and specifically whether this event leads to eventual neuronal death remain undefined. While this presents an intriguing mechanism for Aβ-induced synaptic dysfunction and a new role for APP, its therapeutic relevance is not immediately clear. Thus, it will be critical to further explore this pathway and understand the downstream events of Aβ-induced synaptic toxicity and caspase activation.

## DECLARATIONS

### Consent for Publication

Not Applicable

### Availability of data and materials

The datasets used and/or analyzed during the current study are available from the corresponding author on reasonable request.

### Authors’ contributions

BJM, GP, ST, EHK, and RM designed experiments. BJM, GP, ST performed experiments. BJM, GP, and ST analyzed data. BJM, GP and EHK wrote the paper.

### Acknowledgements

We gratefully thank Dr. Hui Zheng for the APP KO and APLP2 KO mice, Sadegh Nabavi for technical assistance and topical insight, Dr. Todd Golde for gift of antibody, and our respective lab members for helpful discussions and support.

